# Machine Learning-Based Protein Microarray Digital Assay Analysis

**DOI:** 10.1101/2020.08.04.236448

**Authors:** Yujing Song, Jingyang Zhao, Tao Cai, Shiuan-Haur Su, Erin Sandford, Christopher Flora, Benjamin H. Singer, Monalisa Ghosh, Sung Won Choi, Muneesh Tewari, Katsuo Kurabayashi

## Abstract

Serial measurement of a large panel of protein biomarkers near the bedside could provide a promising pathway to transform the critical care of acutely ill patients. However, attaining the combination of high sensitivity and multiplexity with a short assay turnaround poses a formidable technological challenge. Here, we developed a rapid, accurate, and highly multiplexed microfluidic digital immunoassay by incorporating machine learning-based autonomous image analysis. The assay achieved 14-plexed biomarker detection at concentrations < 10pg/mL with a sample volume < 10 μL, including all processes from sampling to analyzed data delivery within 30 min, while only requiring a 5-min assay incubation. The assay procedure applied both a spatial-spectral microfluidic encoding scheme and an image data analysis algorithm based on machine learning with a convolutional neural network (CNN) for pre-equilibrated single-molecule protein digital counting. This unique approach remarkably reduced errors facing the high-capacity multiplexing of digital immunoassay at low protein concentrations. Longitudinal data obtained for a panel of 14 serum cytokines in human patients receiving chimeric antigen receptor-T (CAR-T) cell therapy manifested the powerful biomarker profiling capability and great potential of the assay for its translation to near-real-time bedside immune status monitoring.

## Introduction

Over the past few years, the approach of providing personalized treatment for severely ill patients based on their individualized molecular profiles has received considerable attention as a next step to advance critical care medicine (1–3). Progress has been made in identifying predictive and prognostic protein biomarkers in critical care which holds great promise in patient stratification (4, 5), disease monitoring (6, 7), and therapy development (3, 8). For example, Cao et al (9) tested patients infected with the 2019 novel coronavirus (COVID-19) and reported that a panel of eight plasma cytokines showing significantly heightened levels allowed them to distinguish a group of severely ill patients from a group of mildly ill patients. However, even with the discoveries of these biomarkers, the medical community still falls behind with adopting the precision medicine approach to treat life-threatening acute illnesses, such as cytokine release syndrome (CRS), acute respiratory distress syndrome (ARDS), which are frequently associated with the 2019 novel coronavirus (COVID-19), (9, 10) due to the lack of a sensitive molecular profiling tool to quickly guide clinical decisions or interventions with a near-real-time assay turnaround (1). Additionally, to monitor the highly heterogeneous and time-pressing illness conditions, high multiplex capacity is equally important as sensitivity and speed for improving diagnosis and prognosis accuracy with rich, comprehensive information on multiple biomarker profiles (1, 9, 11, 12). Currently, the commonly used clinical tools for multiplex serum/plasma protein analysis (13), including the bead-based assay coupled flow cytometry and the western blot, fall short of achieving the performance needed for critical care as they require a long assay time (>4 hrs), and laborious steps with limited sensitivity.

Researchers have developed rapid (14–17), point-of-care (18–20), multiplex (21, 22) immunoassays powered by microfluidics. Nonetheless, it is still challenging for these assays to simultaneously achieve a combination of high multiplexity and sensitivity with a rapid assay turnaround time in a clinical setting. By counting single-molecule reactions in fL-nL-volume microwells or droplets (23–25), digital immunoassays can provide unprecedented sensitivity (sub-fM detection) for biomarker analysis. Our recent study (26) demonstrated that it is feasible to extend the digital assay approach to achieve near-real-time protein biomarker profiling at a clinically relevant pM-nM range by quenching reagent reaction at an early pre-equilibrium stage with an incubation process as short as 15-300 sec. However, existing digital immunoassay platforms (27) have limited multiplex capacity (up to 6-plex). The current method (24, 28) utilizes fluorescence dye-encoded beads to identify different analytes. Unfortunately, the nature of binary-based statistical counting brings a few critical challenges to multiplexing digital immunoassays with this method. First, the assay typically requires a large number of beads (e.g. Simoa uses 100,000 beads per plex (27)) for reliable analyte quantification. Mixing and counting such a large number of multi-color-encoded beads tends to cause false signal recognition due to optical crosstalk or non-uniform color coating. Second, increasing multiplexity while keeping the assay’s sensitivity and accuracy additionally requires a large number of microwell arrays to accommodate the large number of beads. This becomes impractical with the current platform as it demands a significantly increased assay device footprint and an image area size. Third, the assay also encounters a significant bead loss during the digitization process partitioning the beads into sub-volumes after the initial reaction process performed for bulk reagent volume in a cuvette (100 μL). All of these issues prohibit the translation of a cheap, robust, point-of-care multiplexed digital assay platform into near-patient applications, thus necessitating a new strategy.

Here, we have developed a highly multiplexed digital immunoassay platform, termed the “pre-equilibrium digital enzyme-linked immunosorbent assay (PEdELISA) microarray.” The PEdELISA microarray analysis integrates on-chip biosensors with a small footprint to minimize the number of images that are needed to read and fully automates the signal counting process, both of which are critically necessary for overcoming the bottlenecks against multiplexing digital assays. The analysis incorporates a powerful microfluidic spatial-spectral encoding method and a machine learning-based image processing algorithm into multi-biomarker detection. The spatial-spectral encoding method confines color-encoded magnetic beads into the arrayed patterns of microwells on a microfluidic chip. The locations of the microwell patterns on the chip indicate which target analytes are detected by trapped color-coded beads. In contrast to the existing digital immunoassay protocol, the fully integrated microfluidic architecture allows the assay reaction to be performed entirely on-chip (no bead loss) at an early pre-equilibrium state, which only requires a sample volume < 10 μL, a 5-min assay incubation and a 75 mm × 50 mm chip size. Based on a convolutional neural network (CNN), the machine learning algorithm permits unsupervised image data analysis while resolving false signal recognition accompanying the multiplexing of digital immunoassays. Employing these biosensing schemes, the PEdELISA microarray platform allows us to simultaneously quantify a large panel of biomarkers in half an hour without sacrificing the accuracy. We used the platform to obtain longitudinal data for blood samples from human patients experiencing cytokine release syndrome (CRS) after chimeric antigen receptor T cell (CAR-T) therapy. The data signify the time-course evolution of the profiles of 14 circulating cytokines over illness development. With its near-real-time assay turnaround and analytical power, the platform manifests great potential to enable acute immune disorder monitoring that guides timely therapeutic interventions.

## Results

### Multiplexed Digital Immunoassay with CNN Image Processing

The PEdELISA microarray analysis used a microfluidic chip fabricated using polydimethylsiloxane (PDMS)-based soft lithography. The chip contains parallel sample detection channels (10–16) on a glass substrate, each with an array of hexagonal biosensing patterns (**Figure 1A**). The hexagonal shape allows each biosensing pattern to densely pack 43,561 fL-sized microwells, which fits into the entire field of view of a full-frame CMOS sensor through a 10x objective lens (**Figure S1)**. Prior to the assay, we deposited magnetic beads (d = 2.8 μm) encoded with non-fluorescent color (no color) and those with Alexa Fluor® 488 (AF 488) into physically separated microwell arrays (**Figure S2**). These beads were conjugated with different capture antibodies according to their colors and locations on the chip. In the current design, the arrangement of 2 colors and 8 arrayed biosensing patterns in each detection channel allows the PEdELISA microarray chip to detect 2 × 8 = 16 protein species (16-plex) at its maximum capacity for each sample loaded to the detection channel. Compared with a single color-encoded method, this combination greatly reduces potential optical crosstalk and fluorescence overlap during a signal readout process. The pre-deposition ensures a fixed number of beads to target each biomarker, which allows more accurate digital counting for each biomarker. It also eliminates bead loss during the conventional partition process and achieves nearly a 100% yield in the signal readout for enzyme active QuantaRed™ (Qred)-emitting beads (“On” beads or “Qred+” beads). The microwell structure (diameter: 3.4 μm and depth: 3.6 μm) was designed to generate sufficient surface tension to hold beads in the microwells. This kept false signals resulting from physical crosstalk between the trapped beads at a negligible level (**Figure S3**).

**Figure 1.**
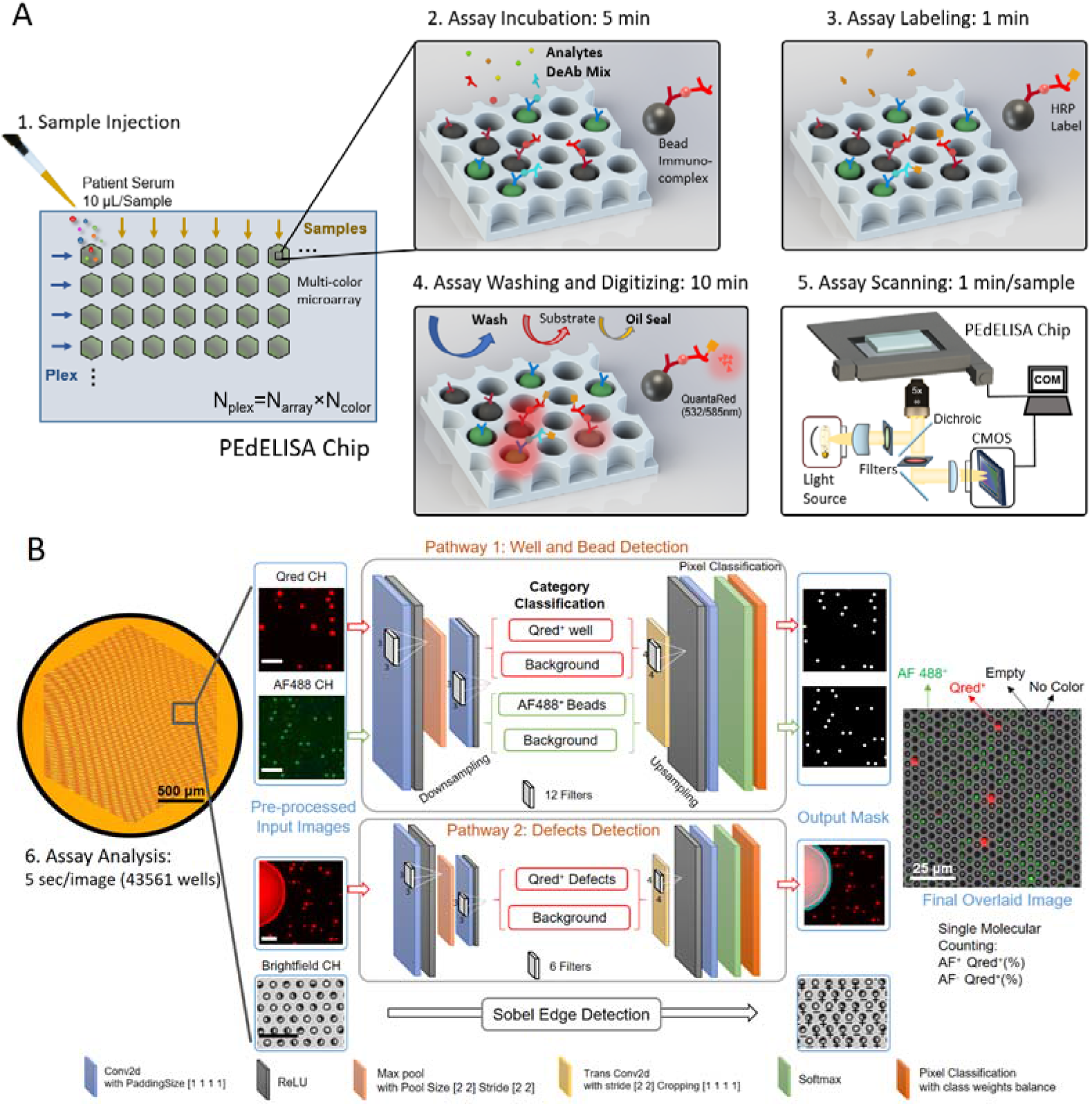
Concept of CNN processed PEdELISA microarray analysis. (A) Microfluidic spatial-spectral encoding method used for multiplexing digital immunoassay. Fluorescence color-encoded magnetic beads coated with different capture antibodies are pre-deposited into the array of hexagonal-shaped biosensing patterns in the microfluidic detection channel. The locations of the biosensing patterns are physically separated from each other. This arrangement yields N_color_× N_array_ measurement combinations determining the assay plexity, N_plex_, where N_color_ is the total number of colors used for encoding beads deposited in each biosensing pattern, and N_array_ is the total number of the arrayed biosensing patterns in each detection channel. In this study, N_color_=2 (non-fluorescent and Alexa Fluor^®^ 488: AF488) and N_array_=8. (B) A convolutional neural network-guided image processing algorithm for high throughput and accurate single molecule counting. Two neural networks were run in parallel, reading multi-color fluorescence image data, recognizing target features versus defects, and generating an output mask for post data processing. The brightfield image was analyzed using a Sobel edge detection algorithm. The images were finally overlaid to determine the fraction of enzyme active beads emitting QuantaRed™ signal (Qred+ beads) to total beads for each color label. The unlabeled scale bars are 25 μm.

A unique challenge posed by highly multiplexed digital assays is to provide fast and accurate analysis of fluorescence signals originating from ~7 million microwells per chip. Additionally, the signal counting process needs to distinguish precisely between images of multi-color bead-filled and empty microwells and to identify signals accurately while subjected to a large fluorescence intensity variance, occasional image defects due to reagent mishandling, and image focus shifts. These challenges make the conventional image processing method with the thresholding and segmentation (GTS) scheme (**Figure S4A**) inaccurate, thus requiring human supervision for error correction in handling digital assay images. Mu et al. (29, 30) showed that the use of machine learning algorithms would provide promising solutions to significantly improve the accuracy of digital assay image processing. However, this approach is only applied for single-color images with a small number of microreactors (a few thousand) with 1080×1120 pixels, which is impractical for high-throughput analysis. To address these challenges, we developed a novel dual-pathway parallel-computing algorithm based on convolutional neural network (CNN) visualization for image processing.

The CNN-based analysis procedure (**Figure 1B**) includes multi-color fluorescence image data read-in/pre-processing (image crop, noise filtering, and contrast enhancement), microwell/bead image segmentation by pre-trained dual-pathway CNN, post-processing, and result output. The key component, dual-pathway CNN, was pre-trained to classify and segment image pixels by labels of (I) fluorescence “On” (Qred channel) microwells, (II) Alexa Fluor® 488 color-encoded beads (AF488 channel), (III) image defects, and (IV) background. The architecture of the network (**Figure 1B**) is separated into a downsampling process for category classification and an upsampling process for pixel segmentation. The downsampling process consists of 3 layers, including 2 convolution layers (4-6 filters, kernel of 3×3) with a rectified linear unit (ReLU), and a max-pooling layer (stride of 2) in between. The upsampling process consists of a transposed convolution layer with ReLU, a softmax layer and a pixel classification layer. To speed up the training process, we started with dividing an image with 32 × 32 pixels and classifying them with the labels and then eventually expanding the image pixel size to 256 × 256 using a pre-trained network (**Figure S5**). We found a large intensity variance across the optical signals from beads in different microwell reactors. As a result, the intensity-based labeling of microwells leads to recognition errors. Microwells with bright beads can be misrecognized to have larger areas with more pixels than those with dim beads. Instead, given that all microwells are lithographically patterned to have an identical size, we labeled them using the same pre-fixed area scale (octagon, r=3 pixel for microwell, disk, r=2 pixel for bead) regardless of their image brightness to make the machine to recognize them correctly. The majority of pixel labels are for the background (Label IV) with no assay information in typical digital assay images. We used the inverse frequency weighting method to further enhance the classification accuracy, which gives more weights to less frequently appearing classes that are identified by Labels (I), (II), and (III) (See **Supporting Information** and **Figure S5** for training details).

In contrast to a previously reported study (29), we greatly reduced the number of convolution layers and filters (depth of network) for high speed processing. Our algorithm employs much fewer labels and features required for imaging processing than those for other typical CNN applications, such as autonomous driving. The unique feature of our algorithm is the ability to run two neural networks in parallel for two detection pathways: one for assay targets (e.g. microwells, beads, and fluorescence signals) and the other for defects. This allows the imaging processing to achieve high speed while maintaining good precision. As a result, it only took ~5 seconds (CPU: Intel Core i7-8700, GPU: NVIDIA Quadro P1000) to process two-color channel data for two 6000×4000 pixel images which contain 43561 micro-reactors.

### PEdELISA Microarray Platform Performance

To validate the effectiveness of the dual-pathway CNN method developed in this work, we compared its performance with that of the standard method based on global thresholding and segmentation (GTS). **Figure 2A** shows representative two-color-channel images causing errors to the image labeling and signal counting of the GTS method. These errors are corrected by the CNN method. For example, false signal counting derives from chip defects or poor labeling reagent confinements within individual microwell reactors due to the local failure of oil sealing. Defocusing can cause two neighboring microwells to be dilated with each other. Highly bright Qred fluorescence from an “On” microwell can cause secondary illumination to light up neighboring microwells. This results in “optical crosstalk” between neighboring microwells (28), which causes the false counting of secondarily illuminated microwells as “On” sites. The uneven illumination of excitation light causes the failure of recognizing dim AF-488 encoded beads (recognized as non-color beads).

**Figure 2.**
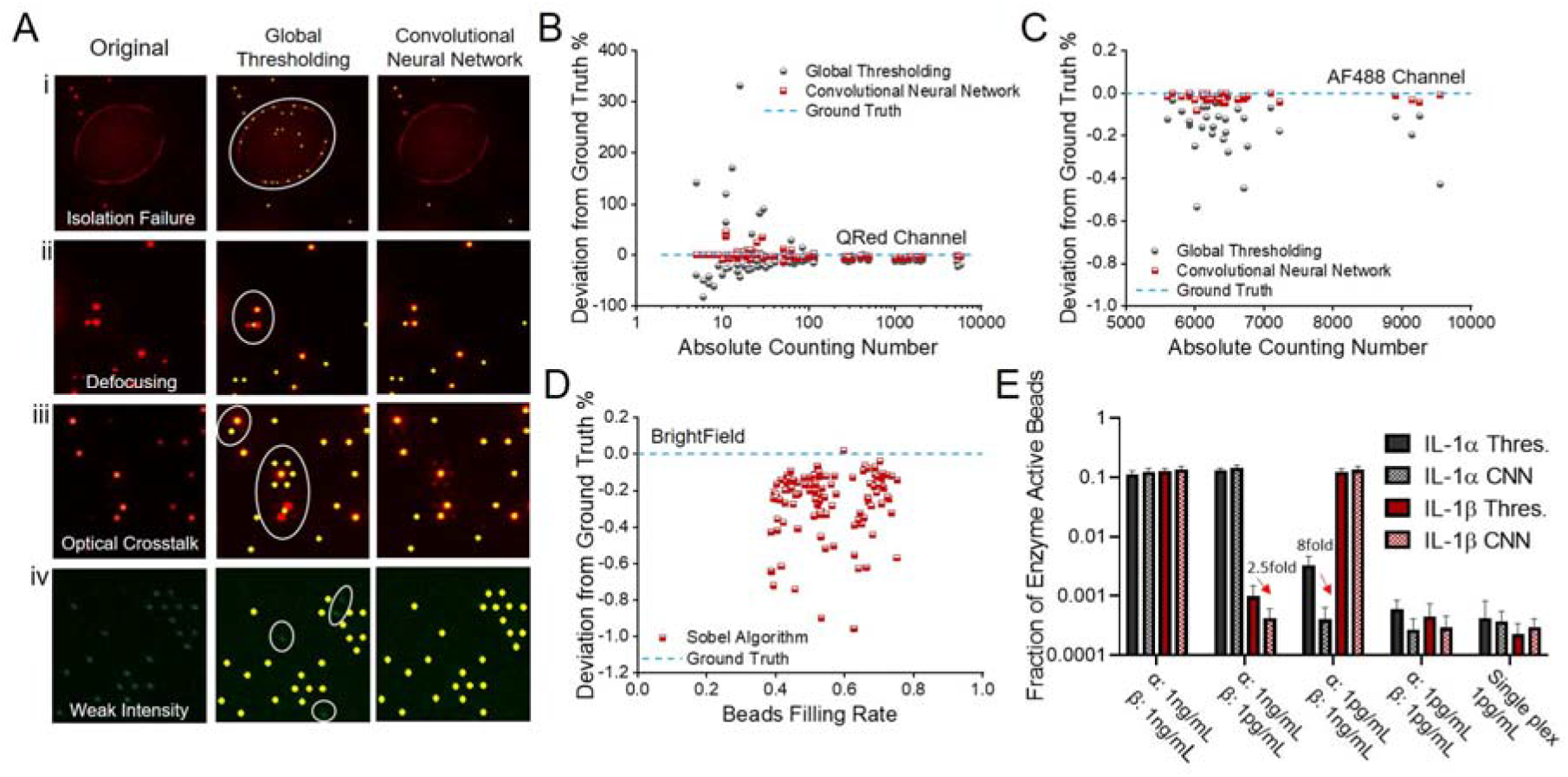
Image processing by convolutional neural network (CNN) and global thresholding and segmentation (GTS) methods (A) representative images causing false signal counting (red dot: Qred+ microwell, green dot: AF-488-colored bead, yellow dot: recognized spot to be counted). (i) The circle represents an area covered by an aqueous reagent solution that is spread over multiple microwell sites due to poor confinement during the oil sealing process. GTS counts potentially false and unreliable signal spots from the area. CNN removes the area from counting. (ii) Image defocusing causes GTS to merge two signal spots from a pair of the neighboring Qred+ microwells in the circle and to count it as a single signal spot. (iii) Secondary illumination of microwell sites due to optical crosstalk in the circle results in their false counting by GTS. (iv) GTS fails to label and count microwell sites holding dim AF-488-colored beads. Error analysis of CNN and GTS methods on (B) Qred-channel (C) AF488-channel and (D) brightfield images. (E) Tests assessing the impact of optical crosstalk on the accuracy of CNN and GTS using dual-color IL-1α and IL-1β detection by spiking (i) IL-1α:1ng/mL IL-1β:1ng/mL (ii) IL-1α:1ng/mL IL-1β:1pg/mL (iii) IL-1α:1pg/mL IL-1β:1ng/mL (iv) IL-1α:1pg/mL IL-1β:1pg/mL (v) IL-1α:1pg/mL IL-1β:1pg/mL assay in single plex for validation. All assays were performed in 25% fetal bovine serum buffer.

In the CNN training process, we collected a large number of images for each error source and used them to train the neural network to achieve results similar to those from manual counting with the human eyes. We applied the following equation to evaluate the error in terms of deviation to the ground truth (%):

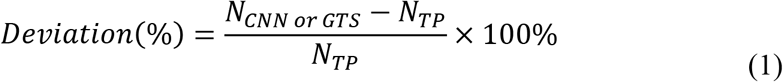

where N_CNN or GTS_ is the number of microwell or bead counted either by CNN or GTS method respectively, N_TP_ is the number of true positives determined by human labeling. The global threshold value was adjusted based on the gray histogram of the image **(Figure S4)**. The human labeling process includes the pre-processing with the GTS method together with human correction to obtain the ground truth, which has been validated by the conventional sandwich ELISA method in our previous study (**Figure S6**) (26).

In counting enzyme active microwells with the Qred channel, we observed that the deviation percentage from ground truth varied with the number of the counted “On” (Qred+) microwells, which is proportional to the analyte concentration. Each data point in **Figure 2B-D** was taken for a hexagonal-shaped biosensing pattern (**Figure 1B**) that contains 43561 microwell arrays with an average bead filling rate of 55.1%. In these data, the number of Qred+ microwells ranged from 1 to 10000 (**Figure 2B**). At higher analyte concentrations (N_Qred_>100), both of the methods achieved reasonably high accuracy with a deviation to the ground truth of 3.92% (CNN) and 9.96% (GTS). However, at the lower concentrations (N_Qred_<100), this deviation became significant (CNN: 5.14% GTS: 71.6%). The larger error of the GTS scheme is attributed to the false counting of regions contaminated with fluorescent reagents and the miscounting of Qred+ microwells of low fluorescence intensity. Thus, the dual-pathway CNN greatly improved the accuracy of the PEdELISA image processing and replacing the thresholding method with our CNN method eliminated the need for human supervision to correct the significant errors in the low concentration region.

In counting color-encoded magnetic beads with the AF-488 channel, we found that the deviation was very small (CNN: 0.021%, GTS: 0.161%). The deviation was suppressed by the little spectral overlap between AF488 and Qred, channels and the high image contrast that we intentionally created between AF-488 and non-color encoded beads (**Figure 2C**). Some miscounting under the uneven spatial distribution of illumination light intensity and the spherical aberration of objective lens over the entire field of view still contributed to the deviation. The CNN method achieved a nearly 8-fold improvement of accuracy. Counting the total number of beads (both no color and fluorescence color-encoded ones) with brightfield images using a customized Sobel edge detection algorithm yielded an average deviation to the ground truth of 0.256% as shown in **Figure 2D**.

To verify our ability to suppress optical crosstalk in the multiplexed assay incorporating the CNN method, we prepared a 25% fetal bovine serum (FBS) sample spiked with two different cytokine species of 1000-fold concentration difference: IL-1α (AF488 encoded) and IL-1β (non-color encoded). Optical crosstalk becomes problematic especially in multiplexed analysis, where the quantity of one biomarker can be serval orders of magnitude higher than those of other biomarkers in the same sample. A slightly false recognition can even give a significantly higher value of biomarker concentration than its true value. **Figure 2E** shows the comparison between the conventional GTS method and the CNN method. False recognition was greatly reduced by the CNN method and we verified that 1pg/mL of IL-1α or β will not interfere even when the other protein reaches 1ng/mL. Furthermore, we performed single-plexed measurements of 1pg/mL of IL-1α and IL-1β, which give “true” concentration values while eliminating optical crosstalk. The single- and dual-plexed measurements both yielded statistically similar results with the CNN method (two-tailed unequal variance t-test, IL-1α P=0.253; IL-1β=0.368), which proves the accuracy of this method even at the presence of strong optical crosstalk.

### Multiplex Pre-Equilibrium Cytokine Detection

Using 2-color encoded (AF488, non-color) magnetic beads with 8 physically separated microarrays, we designed a microfluidic chip to detect 14 cytokines (up to 16-plex) simultaneously (see chip design in Figure S7). Figure 3A shows standard curves obtained from PEdELISA microarray analysis with CNN image processing for cytokines ranging from 0.16 pg/mL to 2.5 ng/mL in 25% FBS. Here, the measurement output is the fraction of the number of enzyme active (Qred+) beads to the total number of beads used for assaying the particular analyte. This fraction is directly correlated to the analyte concentration. The assay was performed for a system at the early state of a transient sandwich immune-complex formation reaction process with a 5-min incubation period, followed by a 1-min enzymatic labeling process. The reaction conditions have been optimized to match all cytokine biomarkers within the clinically relevant range, and a linear dynamic range of three orders of magnitude was achieved in general. Table S2 summarizes the values of the limit of detection (LOD) and limit of quantification (LOQ) of the assay for each cytokine. The antibody-antigen affinity affects the LOD of the assay, and it varies across the detected cytokine species. As a result, we obtained different LOD values for these cytokines even if the capture antibody-conjugated beads were prepared by the same protocol regardless of the analyte types (See Experimental Section). The LOD value tends to decrease with an increasing incubation period. Although the assay was performed with a short incubation period of 5 min, the LOD was found to be still below 5pg/mL (with IL-1β reaching the lowest 0.188pg/mL) after optimizing the detection antibody mixing ratio and the enzyme labeling concentration.

**Figure 3.**
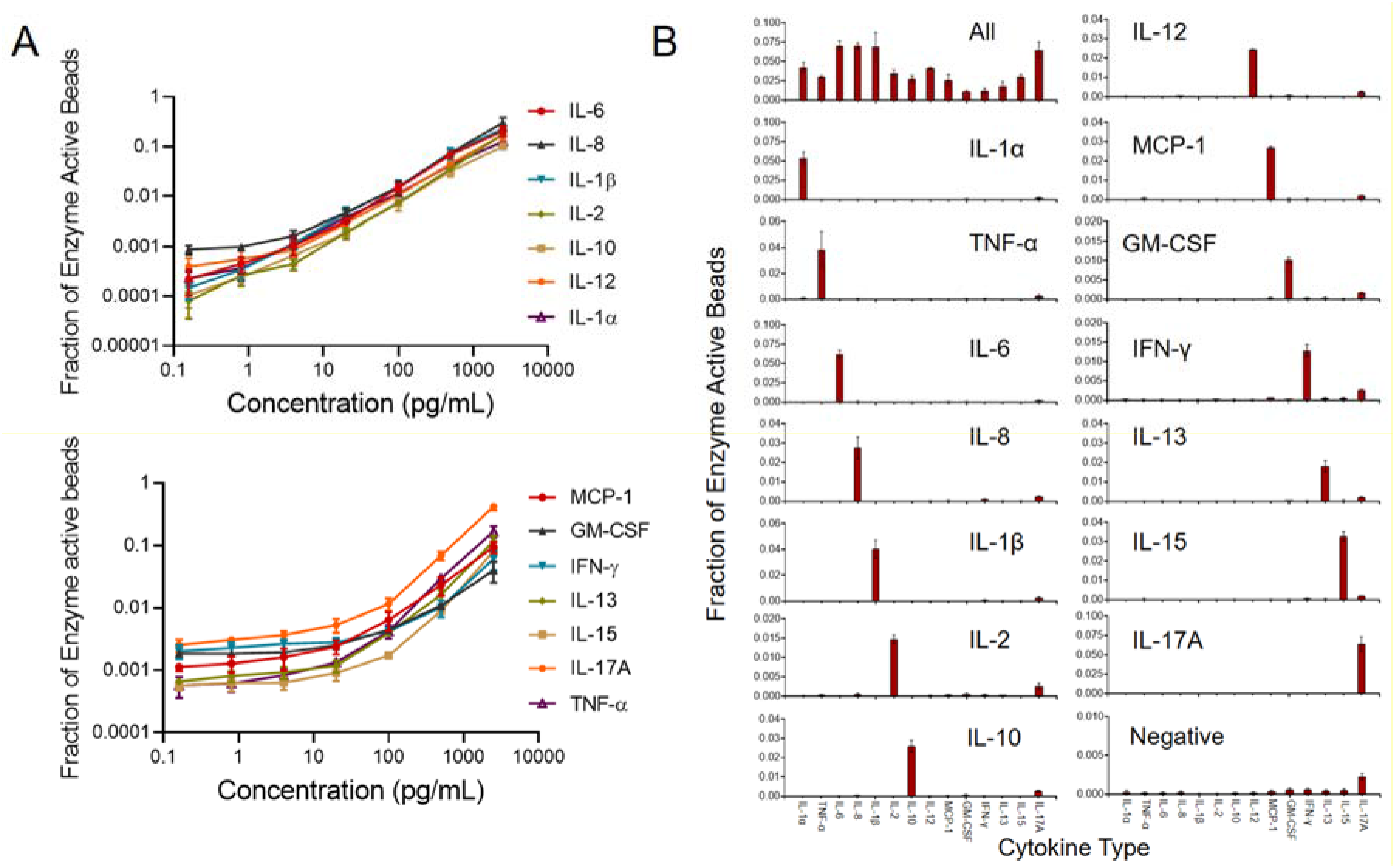
PEdELISA microarray analysis. (A) Assay standard curves for 14 cytokines from 0.16pg/mL to 2500pg/mL in 25% fetal bovine serum (FBS). (B) Assay specificity test with 25% FBS “all-spike-in,” “single-spike-in,” and “no-spike-in” (negative) samples. The analyte concentration of 500pg/mL used for spiking FBS is the optimal value to assess both false positive and negative signals.

We further assessed the level of antibody cross-reactivity among 14 cytokines in 25% FBS. **Figure 3B** shows the assay results for sera spiked by all, one, or none of the recombinant cytokines of 14 species, namely “all-spike-in,” “single-spike-in,” and “no-spike-in” samples. We observed more than 100 times lower background signals from the no-spike-in (negative) sample than those from the all-spike-in sample across the 14 cytokines (except IL-17A for which there is a slightly higher background due to the more active binding between its capture and detection antibodies). The signal-level variation across the 14 cytokines at the same concentration from the all-spike-in and single-spike-in samples could derive from the different levels of analyte-antibody affinity for these cytokines. We also observed a similar trend in the variation of the LOD values for the cytokines from the curves in **Figure 3A**. The signal from each of the 14 single-spike-in samples manifests a high level of specificity to the target analyte. This verifies that the multiplexed assay measurements cause negligible cross-reactivity between each cytokine analyte and other capture and detection antibodies that should not couple with it.

Finally, we applied the 14-plex PEdELISA microarray analysis for the longitudinal serum cytokine measurement from patients receiving CAR-T cell therapy. CAR-T therapies have demonstrated remarkable anti-tumor effects for treatment-refractory hematologic malignancies (31, 32). Unfortunately, up to 70% of leukemia and lymphoma patients who receive CAR-T therapy experience cytokine release syndrome (CRS). CRS is a potentially life-threatening condition of immune activation caused by the release of inflammatory cytokines (e.g., IL-6, TNF-α, and others) (33, 34). CRS initially causes fevers and other constitutional symptoms that can rapidly (i.e., within 24 hours) progress to hypotension and organ damage requiring intensive care. Previous studies (11, 12) have shown the measurement of a panel of cytokines can indicate the early onsite of severe CRS. Thus, the way of intervening CRS could be significantly improved by the multiplex PEdELISA microarray analysis.

To demonstrate the clinical utility of the assay technology, we ran our assay for two CAR-T patients, one who experienced up to grade 2 CRS and one who did not experience CRS in the first few days of post CAR-T infusion. The total sample-to-answer time achieved was 30 min for the entire 14-plexed measurement including the sample incubation (5 min), labeling (1 min), washing/reagent confining (10 min), and image scanning/analysis (14 min) processes. **Figure 4A** shows that Patient 1 developed CRS on day 4 that persisted until day 9. We found significant elevations for all assayed cytokines on Day 0 in comparison to their baseline levels on Day −2 and Day −9. Interestingly, the spike on Day 0 is not due to the CAR-T cells, as the blood sample was taken prior to CAR-T infusion. Typically, CRS patients exhibit a high IL-6 concentration within their blood (35). However, Patient 1 manifested a significantly higher level of TNF-α. This suggests biological heterogeneity in the pathogenic cytokine profiles of patients who develop CRS. We also conducted a similar analysis for a patient who did not develop CRS (**Figure 4B**). We recorded an increase in IL-2 and a relatively high level of IL-17A for this patient, while other cytokines showed no significant changes throughout the analysis. Presumably, normal CAR-T cell expansion was taking place in the patient’s body.

**Figure 4.**
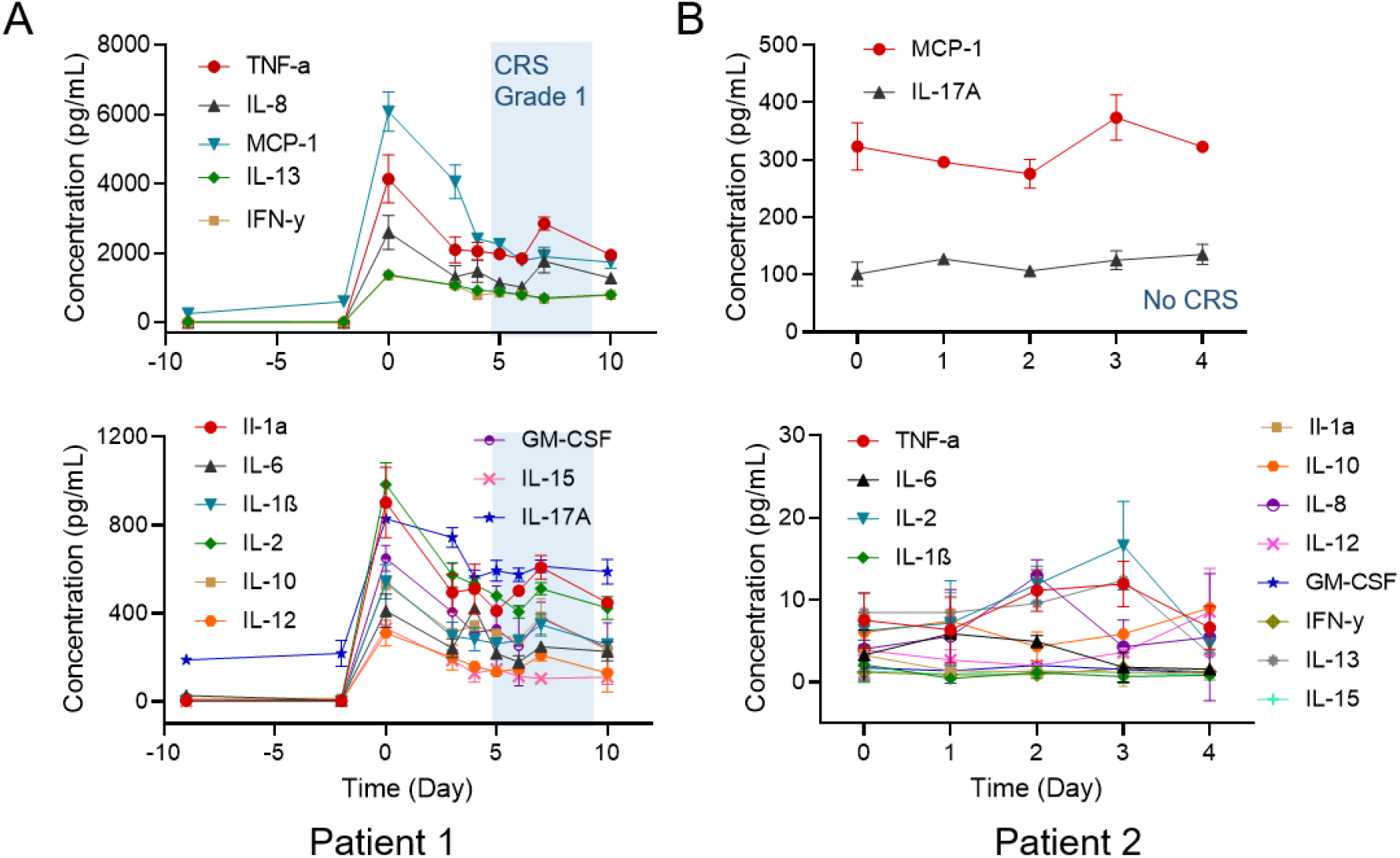
14-plex cytokine measurements in longitudinal serum samples from CAR-T patients who were diagnosed (A) grade 1-2 CRS (B) no CRS. Day 0 represents the day of CAR-T cell infusion. Data before Day 0 represents the baseline. The shaded region marks the period that the patient was diagnosed with grade 1-2 CRS. For better visualization, the data was organized and separately plotted based on the cytokine level from high to low.

## Discussion

We expect that extending the multiplex capacity of digital immunoassay would greatly broaden its utility in the continuous monitoring of protein biomarkers for critically ill patients. However, multiplexing the assay becomes enormously difficult with an increasing number of target biomarkers. Multiplexed digital signal counting required over more than a few millions of fL-sized reactors with conventional methods experiences poor sample/reagent handling and declined accuracy due to various error sources. In this study, we developed a highly multiplexed digital immunoassay platform, namely the PEdELISA microarray, to provide a promising solution for these challenges. The assay platform employs a unique combination of spatial-spectral encoding and machine learning-based image processing on a microfluidic chip. The positional registration of on-chip biosensing patterns, each with more than 40,000 microwell reactors confining sample sub-volumes, fluorescence-encoded analyte-capturing beads, and assay reagents, enabled 14-plex cytokine detection for 10 ◻L of serum with high sample handling efficiency, small reagent loss, and negligible sensor cross talk. The signal processing and analysis of the 14-plexed PEdELISA microarray analysis employed a novel parallel computing CNN-based machine-learning algorithm. This algorithm achieved autonomous classification and segmentation of image features (e.g. microwells, beads, defects, backgrounds) at high throughput (1 min/analyte). Notably, it yielded 8-10-fold higher accuracy than the conventional GTS-based algorithm without any human-supervised error correction.

We ran the PEdELISA microarray measurement of human serum samples from patients who received CAR-T cancer therapy with an incubation time as short as 5 min. The assay simultaneously detected 14 cytokine biomarkers per sample with a clinically relevant dynamic range of pM-nM, and the entire assay process from sample loading to data delivery was completed within 30 min. We tested blood samples obtained from a CAR-T patient at different time points during the course of the therapy with the short assay turnaround. The longitudinal measurement proved the ability of our assay platform to continuously monitor a large number of cytokine profiles that were rapidly evolving in the circulatory system of a patient manifesting CRS. With its speed, sensitivity, multiplexing capacity, and sample-sparing capability, the PEdELISA microarray is poised for future translation to critical care medicine, which is expected to allow the treatment of life-threatening illnesses caused by emerging diseases (e.g., COVID-19) to be timely and tailored with the patient’s comprehensive biomarker profiles.

## Materials and Methods

### Materials

We purchased human IL-6, TNF-α, IL-2, IL-8, IL-13 capture, and biotinylated detection antibody pairs from Invitrogen™, and IL-1α, IL-1β, IL-10, IL-12, IL-15, IL-17A, IFN-γ, GM-CSF and MCP-1 from BioLegend. We obtained Dynabeads, 2.7μm-diameter carboxylic acid, and epoxy-linked superparamagnetic beads, avidin-HRP, QuantaRed™ enhanced chemifluorescent HRP substrate, Alexa Fluor™ 488 Hydrazide, EDC (1-ethyl-3-(3-dimethylaminopropyl) carbodiimide hydrochloride), Sulfo-NHS (Sulfo-N-hydroxysulfosuccinimide), MES (2-(N-morpholino)ethanesulfonic acid) buffered saline, bovine serum albumin (BSA), TBS StartingBlock T20 blocking buffer, and PBS SuperBlock blocking buffer from Thermo Fisher Scientific. We obtained Phosphate buffered saline (PBS) from Gibco™, Sylgard™ 184 clear polydimethylsiloxane (PDMS) from Dow Corning, and Fluorocarbon oil (Novec™ 7500) from 3M™.

### Antibody Conjugation to Magnetic Beads

We prepared the non-color encoded magnetic beads by conjugating epoxy-linked Dynabeads with the capture antibody molecules at a mass ratio of 6 μg (antibody): 1 mg (bead). The Alexa Fluor™ 488 (AF488) encoded magnetic beads were prepared by first labeling carboxylic acid-linked Dynabeads with AF 488 Hydrazide dye and then by conjugating the beads with capture antibody at a mass ratio of 12 μg (antibody): 1 mg (bead) using standard EDC/sulfo-NHS chemistry. Detailed protocol has been described in the previous publication (26). We stored the antibody-conjugated magnetic beads at 10 mg beads/mL in PBS (0.05% T20 + 0.1% BSA + 0.01% Sodium Azide) buffer wrapped with an aluminum foil sheet at 4 °C. No significant degradation of these beads was observed within the 3-month usage.

### Patient Blood Sample Collection and Preparation

Blood samples were collected from patients receiving CAR-T cell therapy and was performed with informed consent under the University of Michigan Institutional Review Boards (IRB) protocol HUM00115179/UMCC 2016.051. Venous blood was collected into a vacutainer containing no anticoagulant on-site at the University of Michigan Medical School Hospital and transported it to a biological lab. After allowing the sample to clot for a minimum of 30 minutes at room temperature, we isolated serum by centrifuging the vacutainer at 1200 × g, for 15 minutes at room temperature. The serum was removed by a pipette, aliquoted into screw cap tubes, and then stored at −80 ºC prior to the assay.

### 14-plex PEdELISA Assay

All assay reagents were prepared in 96-well plate low retention tubes and kept on ice until use. The reagent preparation involved preparing a mixture of biotinylated detection antibody (up to 14 cytokines for CAR-T study) in carrier protein buffer (0.1% BSA, 0.02% Sodium Azide) and storing it at 4C, and preparing an Avidin-HRP solution in a superblock buffer at 100 pM. For the PEdELISA chip calibration, we prepared a mixture of recombinant proteins in 25% fetal bovine serum (standard solution), which was 5x serially diluted from 2.5 ng/mL to 0.16 pg/mL. Prior to the assay, we diluted patient serum samples (5uL) two times with PBS (5uL) to prepare a sample solution. As the first step of the assay, we mixed the sample solution (10 μL) and the biotinylated detection antibody solution (10 μL) (sample mixture) and mixed the 5 titrated standard solutions (10 μL) and the biotinylated detection antibody solution (10 μL) (standard mixtures). Then, we loaded these sample and standard mixtures into the detection channels in parallel and incubated the chip for 300 sec. The signals obtained from the standard mixtures were used for calibrating the biosensors of the chip. The microfluidic channels were then washed with PBS-T (0.1% Tween20) at 20 ◻L/min by s syringe pump for 2 min. 40 μL of the avidin-HRP solution was then loaded into the channel and incubate for 1 min. The chip was washed again with PBS-T (0.1% Tween20) at 20 ◻L/min for 10 min. 30 μL of the enhanced chemifluorescent HRP substrate QuantaRed solution was loaded into the channels and subsequently sealed with 35 μL of fluorinated oil (HFE-7500, 3M). The inlets and outlets of the channels were covered by glass coverslips to prevent evaporation during the imaging process. A programmable motorized fluorescence optical microscopy system was used to scan the image of the bead-filed microwell arrays on the microfluidic chip, identify the bead type (non-color vs. AF488 dyed), and detect the enzyme-substrate reaction activity. This system is composed of a Nikon Ti-S fluorescence microscope (10x objective), a programmable motorized stage (ProScan III), a mercury lamp fluorescence illumination source, a SONY full-frame CMOS camera (α7iii), and a custom machined stage holder. The motorized stage was pre-programmed to follow the designated path to scan the entire chip (160 images) in 3 sequential steps: 1. Scan the QuantaRed channel (532nm/585nm, excitation/emission) 2. Scan the AF488 channel (495nm/519nm, excitation/emission) 3. Scan the brightfield. It typically took around 5-7 min to scan the entire chip for 10 samples in 16-plex detection.

### Statistics

Experiments with both synthetic recombinant proteins and CAR-T patient samples at each time point were performed 3 times (in independent tests) with two on-chip repeats to obtain the error bar. Group differences were tested using a two-tailed unequal variance t-test. A p-value of < 0.05 was considered to be statistically significant.

## Data availability

The CAR-T patient cytokine data in this study is available through the database. All relevant data are available within the article file or Supplementary Information, or available from the authors on reasonable request.

## Acknowledgments

This study was supported by the National Science Foundation (CBET1931905), the U-M Precision Health Scholars Award (Y.S.), the A. Alfred Taubman Medical Research Institute (M.T. and S.W.C.), and the Cancer Research Institute AWD012546 (M.T., S.W.C., and K.K.). The authors also acknowledge clinical support from the Hematology/Oncology Division at the Department of Internal Medicine and the CAR-T patients who participated in this study under the UM IRB protocol HUM00115179/UMCC 2016.051. Device fabrication was performed at the UM Robert H. Lurie Nanofabrication Facility.

## References

1. A. Sarma, C. S. Calfee, L. B. Ware, Biomarkers and Precision Medicine: State of the Art. Crit Care Clin 36, 155–165 (2020).

2. C. W. Seymour et al., Precision medicine for all? Challenges and opportunities for a precision medicine approach to critical illness. Critical Care 21 (2017).

3. T. van der Poll, F. L. van de Veerdonk, B. P. Scicluna, M. G. Netea, The immunopathology of sepsis and potential therapeutic targets. Nat Rev Immunol 17, 407–420 (2017).

4. C. S. Calfee et al., Acute respiratory distress syndrome subphenotypes and differential response to simvastatin: secondary analysis of a randomised controlled trial. Lancet Respiratory Medicine 6, 691–698 (2018).

5. H. R. Wong et al., Interleukin-8 as a stratification tool for interventional trials involving pediatric septic shock. Am J Resp Crit Care 178, 276–282 (2008).

6. J. D. Faix, Biomarkers of sepsis. Crit Rev Clin Lab Sci 50, 23–36 (2013).

7. S. Kibe, K. Adams, G. Barlow, Diagnostic and prognostic biomarkers of sepsis in critical care. J Antimicrob Chemother 66 Suppl 2, ii33–40 (2011).

8. P. Schuetz et al., Effect of procalcitonin-guided antibiotic treatment on mortality in acute respiratory infections: a patient level meta-analysis. Lancet Infect Dis 18, 95–107 (2018).

9. C. Huang et al., Clinical features of patients infected with 2019 novel coronavirus in Wuhan, China. Lancet 395, 497–506 (2020).

10. N. Chen et al., Epidemiological and clinical characteristics of 99 cases of 2019 novel coronavirus pneumonia in Wuhan, China: a descriptive study. Lancet 395, 507–513 (2020).

11. K. A. Hay et al., Kinetics and biomarkers of severe cytokine release syndrome after CD19 chimeric antigen receptor-modified T-cell therapy. Blood 130, 2295–2306 (2017).

12. D. T. Teachey et al., Identification of Predictive Biomarkers for Cytokine Release Syndrome after Chimeric Antigen Receptor T-cell Therapy for Acute Lymphoblastic Leukemia. Cancer Discovery 6, 664–679 (2016).

13. L. Cohen, D. R. Walt, Highly Sensitive and Multiplexed Protein Measurements. Chem Rev 119, 293–321 (2019).

14. Y. J. Song et al., AC Electroosmosis-Enhanced Nanoplasmofluidic Detection of Ultralow-Concentration Cytokine. Nano Lett 17, 2374–2380 (2017).

15. X. T. Tan et al., Glass capillary based microfluidic ELISA for rapid diagnostics. Analyst 142, 2378–2385 (2017).

16. W. W. Jing et al., Time-Resolved Digital Immunoassay for Rapid and Sensitive Quantitation of Procalcitonin with Plasmonic Imaging. Acs Nano 13, 8609–8617 (2019).

17. Y. Park et al., An Integrated Plasmo-Photoelectronic Nanostructure Biosensor Detects an Infection Biomarker Accompanying Cell Death in Neutrophils. Small 16 (2020).

18. B. Reddy et al., Point-of-care sensors for the management of sepsis. Nat Biomed Eng 2, 640–648 (2018).

19. Y. Park et al., Biotunable Nanoplasmonic Filter on Few-Layer MoS2 for Rapid and Highly Sensitive Cytokine Optoelectronic lmmunosensing. Acs Nano 11, 5697–5705 (2017).

20. J. Min et al., Integrated Biosensor for Rapid and Point-of-Care-Sepsis Diagnosis. Acs Nano 12, 3378–3384 (2018).

21. P. Y. Chen et al., Multiplex Serum Cytokine Immunoassay Using Nanoplasmonic Biosensor Microarrays. Acs Nano 9, 4173–4181 (2015).

22. R. Fan et al., Integrated barcode chips for rapid, multiplexed analysis of proteins in microliter quantities of blood. Nat Biotechnol 26, 1373–1378 (2008).

23. D. M. Rissin et al., Single-molecule enzyme-linked immunosorbent assay detects serum proteins at subfemtomolar concentrations. Nat Biotechnol 28, 595–599 (2010).

24. V. Yelleswarapu et al., Mobile platform for rapid sub-picogram-per-milliliter, multiplexed, digital droplet detection of proteins. Proc Natl Acad Sci U S A 10.1073/pnas.1814110116 (2019).

25. Y. Zhang, H. Noji, Digital Bioassays: Theory, Applications, and Perspectives (vol 89, pg 92, 2017). Anal Chem 89, 13675–13675 (2017).

26. Y. J. Song et al. (2020) Rapid single-molecule digital detection of protein biomarkers for near-real-time monitoring of systemic immune disorders. in Submitted to Blood, p Under Review.

27. A. J. Rivnak et al., A fully-automated, six-plex single molecule immunoassay for measuring cytokines in blood. J Immunol Methods 424, 20–27 (2015).

28. D. M. Rissin et al., Multiplexed single molecule immunoassays. Lab Chip 13, 2902–2911 (2013).

29. Z. M. Hu et al., A novel method based on a Mask R-CNN model for processing dPCR images. Anal Methods-Uk 11, 3410–3418 (2019).

30. T. Gou et al., A new method using machine learning for automated image analysis applied to chip-based digital assays. Analyst 144, 3274–3281 (2019).

31. S. L. Maude et al., Tisagenlecleucel in Children and Young Adults with B-Cell Lymphoblastic Leukemia. N Engl J Med 378, 439–448 (2018).

32. S. S. Neelapu et al., Axicabtagene Ciloleucel CAR T-Cell Therapy in Refractory Large B-Cell Lymphoma. N Engl J Med 377, 2531–2544 (2017).

33. S. S. Neelapu et al., Chimeric antigen receptor T-cell therapy - assessment and management of toxicities. Nature Reviews Clinical Oncology 15, 47–62 (2018).

34. D. W. Lee et al., Current concepts in the diagnosis and management of cytokine release syndrome. Blood 124, 188–195 (2014).

35. C. Kotch, D. Barrett, D. T. Teachey, Tocilizumab for the treatment of chimeric antigen receptor T cell-induced cytokine release syndrome. Expert Rev Clin Immunol 15, 813–822 (2019).

