## Supporting Information for "Machine Learning-Based Protein Microarray Digital Assay Analysis"

* Katsuo Kurabayashi

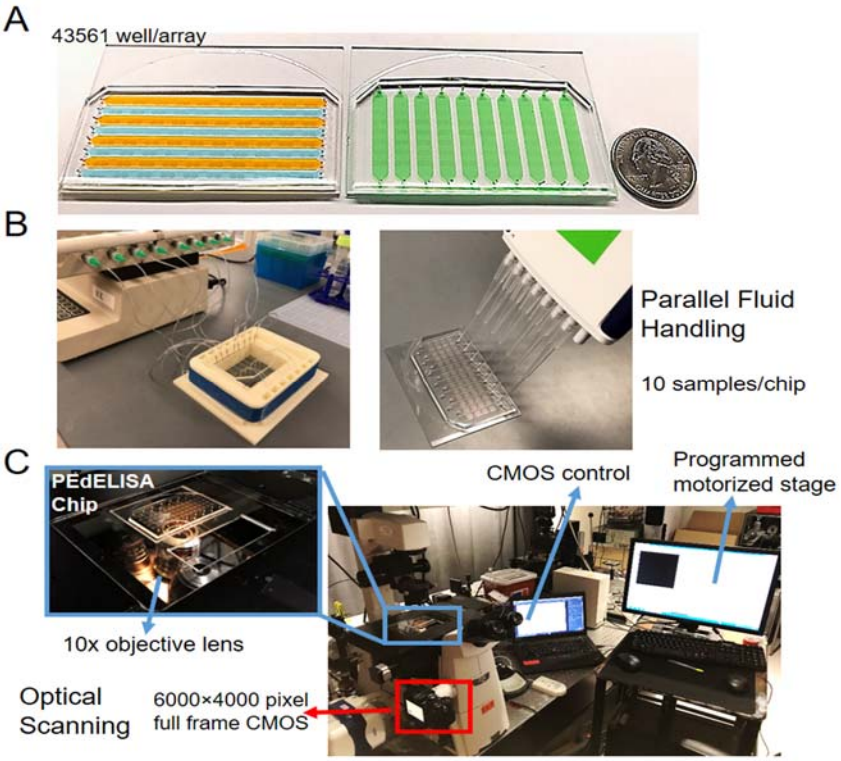


**Figure S1.** The PEdELISA immunoassay system employing (A) a PEdELISA microarray chip prepared with a multi-array biosensor layer attached to a bead patterning layer (left) and to a sample loading layer (right), (B) a parallel fluid handling unit using a multichannel pipette for sample loading and a syringe pump for on-chip washing, and (C) a programmed dual-color fluorescence optical scanning setup.

**Microfluidic Chip Fabrication and Spatial-Spectral Encoding:** The first step of the PEdELISA microarray chip fabrication involved the construction of three different PDMS layers (**Figure S2**). The first PDMS layer has arrays of hexagonal biosensing patterns with microwell (d=3.4 μm) structures (multi-array biosensor layer). The second one has bead settling flow channels of 2500 μm in width, 70 μm in height, and 65 mm in length (bead setting layer). The third one has channels of 4.5 mm in width, 90 μm in height, and 30 mm in length for the detection of analytes in loaded samples (sample detection layer). First, we constructed SU-8 molds for these three PDMS layers on separate oxygen plasma treated silicon wafers by standard photolithography. This process involved depositing negative photoresist (SU-8 2005, SU-8 2050, MicroChem) layers at different spin coating speeds to form the designed thicknesses for the PDMS microstructure patterns. Subsequently, a precursor of PDMS was prepared at a 10:1 base-to-curing-agent ratio. To construct the multi-array biosensor layer, we deposited a thin PDMS precursor film (~300 μm) onto the microwell-patterned SU-8 mold by spin coating, baked it overnight (60 °C), and then attached it to a pre-cleaned 75×50mm glass substrate through oxygen plasma treatment. We made both of the bead settling layer and the sample detection layer by pouring the PDMS precursor over the other SU-8 molds in a petri dish and then baked overnight (60 °C).

The second step involved the settlement of beads in the microwells of each hexagonal patter on the multi-array biosensor layer. We first aligned and attached the bead setting layer to the multi-array biosensor layer on the glass substrate. Then, we prepared 7 sets of a 25 uL mixture of AF488 encoded beads (anti-cytokine 1) and non-color encoded beads (anti-cytokine 2) at the concentration of 1mg/m in vials for the 14-plex detection. This was followed by loading each of the 7 mixtures into one of the microfluidic channels in the bead settling layer (**Figure S2A**). After waiting for the beads to settle in the microwells for 5 min, we washed the bead settling channels with 200 uL PBS-T (0.1% Tween20) to remove the unstrapped beads thoroughly. At this step, we ensured that the microwells were filled with the beads at a sufficient rate (typically above 50%) using an optical microscope. (If not, the bead mixture solution was reloaded and washed again.)

The third step involved the assembly of the chip with the multi-array biosensor and sample detection layers. After the bead setting channels were dried by sucking out the washing buffer using a pipette, we peeled off the bead settling layer from the multi-array biosensor layer and replaced it with the sample detection layer. Here, the sample detection layer was aligned and attached to the multi-array biosensor layer so that its channels were oriented perpendicular to the direction of the channels of the bead settling layer. We then slowly loaded the sample detection channels with Superblock buffer (0.05% Tween20) to passivate the PDMS surface and incubate the whole chip for at least 1 hour before the assay to avoid non-specific protein adsorption. The sample detection layer was punched to form inlets and outlets for its channels. The chip was tape cleaned and covered before the assay usage. Finally, serum samples were loaded to the sample detection channels from their inlets (**Figure S2D**).

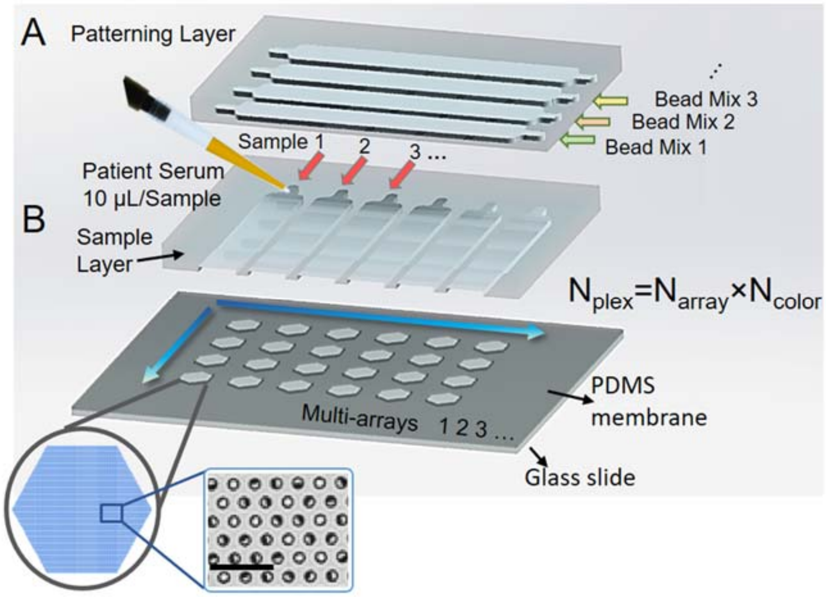


**Figure S2.** Multiplexing the digital immunoassay by constructing a PEdELISA microarray chip based on the concept of microfluidic spatial-spectral encoding: (A) Trapping beads into microwells of the arrayed biosensing patterns on the multi-array biosensor layer. Mixtures of fluorescently encoded beads of $N_{color}$ colors are loaded to the microfluidic channels on the bead settling layer. (B) Peeling off the bead settling layer from the multi-array biosensor layer, and attaching the sample loading layer on the multi-array biosensor layer. The 90º orientation of the sample loading channels permits each channel to contain an array of biosensing patters of $N_{array}$ types. Loading serum samples to the channels of the sample detection layer with pipettes. The chip arrangement yields a total of $N_{color}$×$N_{array}$ plex for the analysis of each sample.

**Bead flushing test for assessing physical crosstalk**: We considered false signals resulting from misplacing beads during the preparation of the PEdELISA chip. If some of the beads targeting analyte *A* were accidentally trapped into the microwell arrays of a biosensing pattern to detect analyte *B*, these misplaced beads would yield either false positive or false negative signals of analyte *B*, thus confounding the assay ("physical crosstalk" between beads). Previous studies (1, 2) reveal that choosing an appropriate design for the PDMS microwell allows surface tension to hold firmly a bead trapped in it even when the entire chip is flipped upside down under prolonged sonication. Guided by these studies, we designed the microwell structure to be 3.4 μm in diameter and 3.6 μm in depth to generate sufficient surface tension to hold beads in microwells (using a permanent magnet could further facilitate the seeding and retention of beads (2,3)).

To assess the impact of physical crosstalk on our assay, we ran a control test. The test started with settling AF-488 encoded magnetic beads into one of the sample loading channels of the chip (AF-488 bead channel), and then subsequently settling non-color encoded beads into another channel next to it (non-color bead channel). Then, we washed away the untrapped beads from these channels, peeled off the bead settling layer from the multi-array biosensor layer, replaced it with sample loading layer, and applied harsh flushing first for the AF-488 bead channel and next for the non-color channel at a washing buffer flow rate of 40uL/min for 15 min. Finally, after sealing the microwells in the channels with oil, we took a fluorescence microscopy image of the chip and evaluated the number of AF-488 encoded beads invading the non-color bead channel before and after the flushing process. Across 160 independent microwell sites, we observed physical crosstalk occurring at an average bead misplacement percentage of 0.087% out of the total beads originally settled in the non-color bead channel with a 0.0012% standard deviation (**Figure S3**). Given that digital ELISA typically forms less than 0.1 antibody-antigen-antibody immune-complexes per bead on average, the false positive signal generated by physical crosstalk is nearly an order less than the typical negative control signal (0.0005-0.001 average molecule per bead) due to non-specific adsorption of proteins.

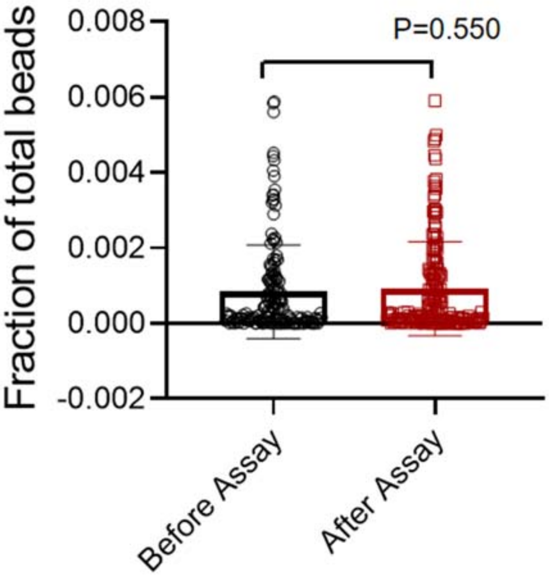


**Figure S3.** Fraction of AF-488 encoded beads invading the channel to be loaded with non-color coded beads before and after the bead flushing test assay (P=0.550). On average, only 0.087% of the trapped beads were misplaced in the channel for the both cases. The result was obtained from the fluorescence images of 160 independent microwell sites. The harsh assay conditions with a washing buffer flow rate of 40uL/min and a duration of 15 min caused negligible physical crosstalk between the beads.

**Global thresholding and segmentation vs convolutional neural network visualization: Figure S4** shows the side-by-side comparison between the global thresholding and segmentation (GTS) method and the convolutional neural network (CNN) method in PEdELISA image processing. Both of the methods start from a pre-processing process, which typically includes image cropping, contrast enhancement and noise filtering. The GTS method involves finding and adjusting a global threshold value based on the gray histogram of the image (Figure S4A, black dash line). This method labels a microwell site showing a fluorescence intensity level above the threshold value as a positive ("On") pixel. Then, it applies a 2×2-pixel image erosion mask along the edge to remove the randomly appearing shot noise pixels with intensities above the threshold. The post-processing process of the GTS method includes image dilation and segmentation, "On" microwell/bead counting, error correction and image overlay (more details can be found in the previous publication (4)).

Whereas, the CNN method runs two signal recognition pathways in parallel, which are pre-trained to recognize enzyme active "On" microwells (Red channel, Qred) or beads (Green channel, AF488) versus defects and contaminations using 5,750 labeled images. As a result, this method does not need to predetermine the intensity threshold value required for the GTS method. The CNN method collects three types of raw images (4000×6000 pixels) for each biosensing pattern with 43,561 microwells according to (1) the red fluorescence channel (Qred), (2) the green fluorescence channel (AF488), and (3) the bright-field channel. The Qred- and AF488-channel images are first cropped and pre-processed for noise filtering and contrast enhancement, and then sent directly to the dual-pathway CNN to recognize the enzyme active Qred fluorescence "On" (Qred+) microwell, AF488 fluorescence emitting (AF488+) bead, image defect, and background image features. Then the Qred+ and AF488+ targets are segmented out as the output mask with the defects are removed. The bright-field image is used to recognize both AF488+ and non-color beads and analyzed using the Sobel edge detection method to determine the overall bead filling rate. After post-image processing, the three images are overlaid to determine the numbers of Qred+ microwells and the bead color types (AF488+ or non-color) within them. The fractional population of the Qred+

microwells with respect to the total bead-filled microwells is determined for both the AF488+ (AF+ Qred+ %) and non-color bead types (AF- Qred+ %) and used to obtain the concentrations of the two different biomarkers. The PEdELISA microarray chip with the current design allows us to analyze up to 16 different analytes for each sample by applying the CNN method-based imaging processing to 8 physically distinct biosensing patterns on it.

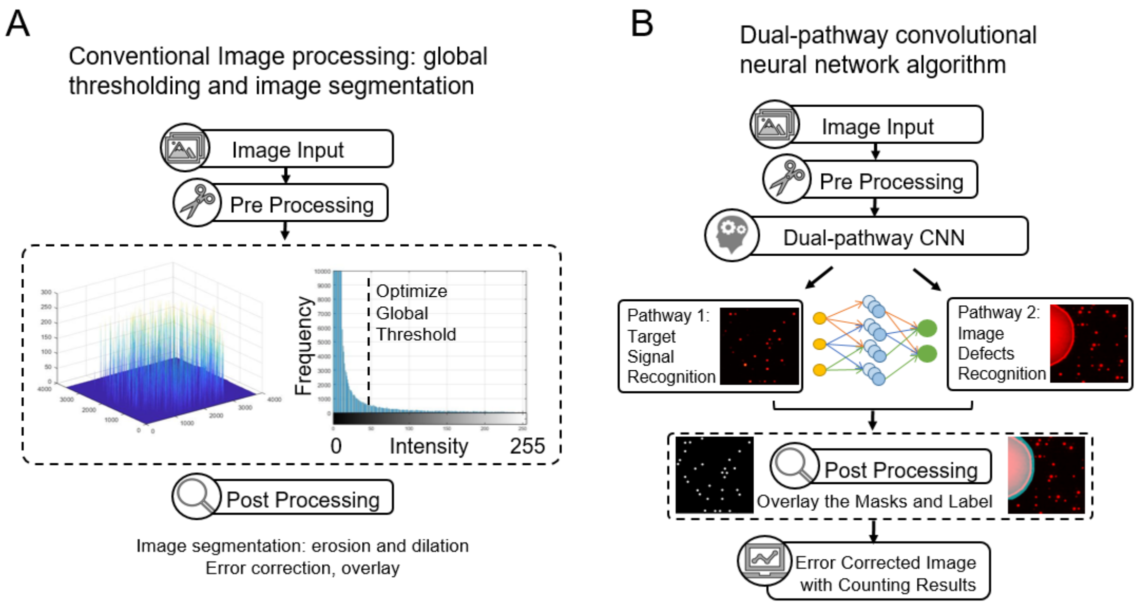


**Figure S4.** Algorithm comparison between the global thresholding segmentation (GTS) and the convolutional neural network (CNN). In GTS, an optimized global threshold value is predetermined based on the intensity histogram of the image to be processed (shown by the black dash line). The CNN method does not require the predetermination of the threshold value.

**Training details of the dual-pathway convolutional neural network: Figure S5** shows the training process of our dual-pathway CNN algorithm, which involves data set preparation and 2-step neural network training. We first developed pre-stage neural networks trained with a data set of 3,000 representative locally-cropped (32×32 pixel) images. For each of these images, two types of images (6,000 in total) only showing the labeling information, which are called "masks" were generated by thresholding and manual labeling (Figure S5A Labeled). In the pre-stage mask for Pathway 1, all the target pixels representing "On" (Qred+) microwell sites and other sites (defects or background) were labeled as '1' and '0', respectively. In the pre-stage mask for Pathway 2, the defects and other sites were labeled as '1' and '0', respectively. The pre-trained networks were able to identify and segment the labeled objects to some degree of accuracy (i.e., 60-80% "On" wells). But they still lacked the accuracy to extract more in-depth features of the object, such as its shape and fluorescence intensity variations of the objects, and other defects that were hard to identify with the small-scale images.

To further improve the accuracy, we built three second-stage parallel CNN networks to identify AF 488+ pixels, Qred+ pixels, and defects that were trained with 256×256-sized images (around 200 images per network). The labeling masks to be used to train the second-stage networks were generated by the pre-stage neural networks with human correction (see Figure S5Bii). By taking into account that the sizes of lithographically patterned microwells and beads were predetermined, we trained the second-stage networks to accurately recognize these objects independent of their image intensity variations (see the main text in Section 2.1). Additionally, we selected 30 images for each error source caused by optical crosstalk (see Figure 2A, ii&iii and main text for detailed definition), carefully determined the intensity threshold to set a clear boundary between adjacent microwells for each of these images, and generated error-free second-stage labeling masks.

As for training the defect recognition-network, we first enhanced the image brightness and contrast to recognize even low-intensity regions. We then applied image dilation for the defect location to ensure the generated mask area is large enough to cover the entire defect area as shown in Figure S5iii. Here, only defect locations were labeled as '1' and all others were labeled as '0'. The region labeled as '1' was eventually removed from the total analyzed area to eliminate it from the fluorescence signal counting.

Our neural network contains 5 classes: Qred+ class, AF488+ class, Qred defect class, Qred background class, and AF488 background class. Before training the neural network with the labeling masks above, we also added weight information to each class to further enhance the pixel identification accuracy. We used the inverse frequency weighting method which gives more weights to less frequently appearing classes. The class weight was defined as

$$\text{Classweight} = \frac{N_{image\ total\ pixels}}{N_{class\ pixels}}$$

where $N_{image\ total\ pixels}$ is the number of total image pixels of 256×256 = 65,536, and the $N_{class\ pixels}$ is the number of pixels for each class. This class weighting strategy was added into the neural network training process because the number of $Qred^+$ or $AF488^+$ class pixels were significantly smaller than the number of their total background pixels.

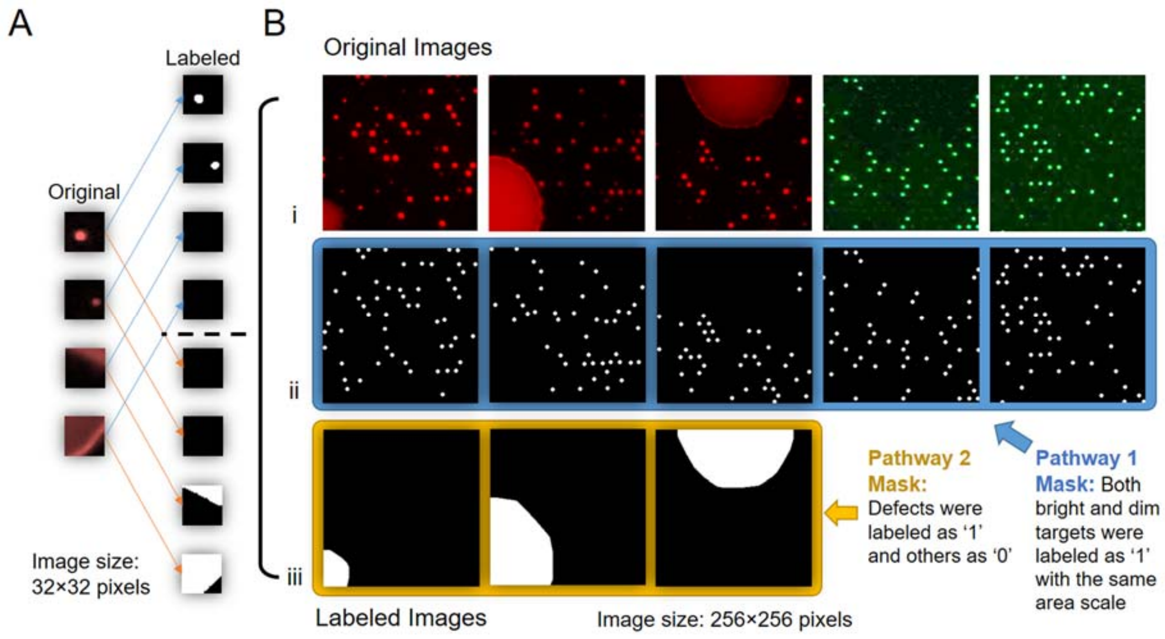


**Figure S5.** Training process of the dual-pathway convolutional neural network with semantic segmentation. (A) The data library was carefully pre-selected based on 3000 representative images (32×32 pixels) to pre-train the CNN. The images were first labeled based on GTS and then manually modified based on human supervision. (B) The pre-trained CNN was then used to label a new data library which contains around 200 larger images (256×256 pixels) to further improve the feature extraction and classification accuracy.

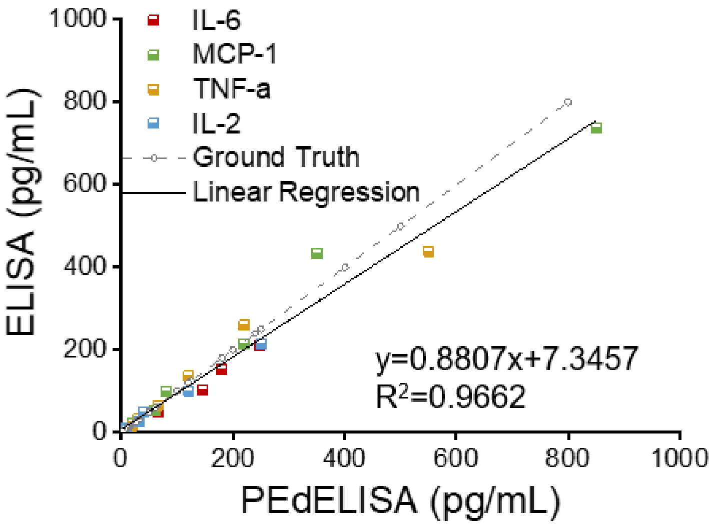


**Figure S6.** Correlation between PEdELISA (5-min incubation) and conventional sandwich ELISA tests for the four selected cytokine IL-6, MCP-1, TNF-α and IL-2 using spike-in recombinant proteins in 25% fetal bovine serum. The ground truth is plotted in dotted line with scattered pre-determined spike-in concentrations. The PEdELISA data was analyzed based on the GTS method with human correction.

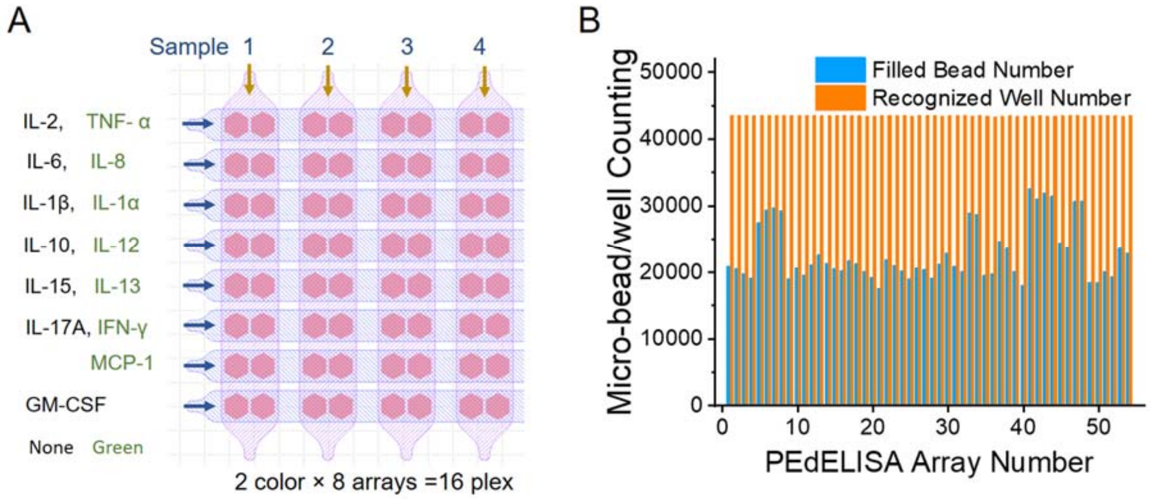


**Figure S7.** (A) Arrangement of a panel of 14 cytokine detection for CAR-T cytokine release syndrome detection test. The two cytokines labeled with the black and green fonts on each row were detected in the sample detection channels (1, 2, 3, and 4) vertical to the row using non-color (black) and AF-488 (green) encoded beads, respectively. (B) Counts of microwells filled with beads and microwells recognized with brightfield images for 60 randomly selected arrayed biosensing patterns (43561 microwells per pattern). The average bead filling rate for the biosensing pattern was 55.1%.

**Table S1.** PEdELISA cost analysis and benchmarking

| Diagnostic System | Cost/96 Assays | Instrument Cost | *Assay Time | LOD (pg/mL) | Sample Volume (µL) | Plexity |
|---|---|---|---|---|---|---|
| Luminex | $3000 | >$40,000 | > 4 hours | 0.1 -100 | 25-50 | 25-65 |
| Colorimetric ELISA | $200 | $5,000 | > 4 hours | 1-10 | 100 | 1 |
| Quanterix SIMOA | $1000 | >$200,000 | 45 min | 0.002 - 1 | 100 | 6 |
| PerkinElmer AlphaLISA | $1200 | >$200,000 | 4 hours | 1-100 | 50-100 | 4 |
| Our PEdELISA Platform | $50 | <$10,000 | < 30 min | 0.1-5 | 5-10 | 16 |

**Table S2.** Limit of detection (LOD), Limit of Quantification (LOQ) and standard root mean square coefficient of variance (RMS CV) summary of 5-min 14-plex PEdELISA. Here, the LOD was determined by concentration from the reagent blank's signal + 3σ and the LOQ was reagent blank's signal + 10σ. The RMS CV was determined by the root mean square average signals from 20, 100, and 500 pg/mL assay standard with typical three day-by-day repeats and 2 on-chip repeats.

| Cytokine Type | TNF-α | IL-6 | IL-8 | IL-1β | IL-2 | IL-10 | IL-12 |
|---|---|---|---|---|---|---|---|
| Assay Blank+3σ (%) | 0.0580 | 0.0456 | 0.0920 | 0.0148 | 0.0222 | 0.0217 | 0.0493 |
| Assay Blank+10σ (%) | 0.1355 | 0.1168 | 0.1820 | 0.0404 | 0.0601 | 0.0584 | 0.1109 |
| LOD 5-min (pg/mL) | 2.195 | 2.667 | 2.033 | 0.188 | 0.535 | 0.210 | 0.613 |
| LOQ 5-min (pg/mL) | 12.80 | 12.21 | 11.54 | 0.696 | 11.24 | 5.661 | 4.996 |
| Assay standard RMS CV (%) | 16.0 | 6.30 | 14.6 | 5.19 | 15.3 | 9.96 | 14.1 |

| Cytokine Type | IL-1α | MCP-1 | IL-13 | IL-15 | IL-17A | IFN-γ | GM-CSF |
|---|---|---|---|---|---|---|---|
| Assay Blank+3σ (%) | 0.0527 | 0.1557 | 0.0706 | 0.0518 | 0.3675 | 0.2283 | 0.2139 |
| Assay Blank+10σ (%) | 0.1240 | 0.3769 | 0.1758 | 0.1299 | 0.6641 | 0.3566 | 0.2896 |
| LOD 5-min (pg/mL) | 0.349 | 3.406 | 1.733 | 3.206 | 4.120 | 3.448 | 7.797 |
| LOQ 5-min (pg/mL) | 2.674 | 47.83 | 18.296 | 57.944 | 43.423 | 62.66 | 35.33 |
| Assay standard RMS CV (%) | 13.1 | 18.8 | 10.1 | 20.6 | 11.1 | 12.6 | 10.7 |

**Reference:**

1. W. H. Henley, P. J. Dennis, J. M. Ramsey, Fabrication of Microfluidic Devices Containing Patterned Microwell Arrays. Anal Chem 84, 1776-1780 (2012).
2. N.-T. Huang, Y.-J. Hwong, R. L. Lai, A microfluidic microwell device for immunomagnetic single-cell trapping. Microfluidics and Nanofluidics 22, 16 (2018).
3. K. Akama et al., Wash- and Amplification-Free Digital Immunoassay Based on Single-Particle Motion Analysis. Acs Nano 13, 13116-13126 (2019).
4. Y. J. Song et al. (2020) Rapid single-molecule digital detection of protein biomarkers for near-real-time monitoring of systemic immune disorders. in Submitted to Blood, p Under Review.
